# The emerging role of physical modeling in the future of structure determination

**DOI:** 10.1101/228247

**Authors:** Kari Gaalswyk, Mir Ishruna Muniyat, Justin L. MacCallum

## Abstract

Biomolecular structure determination has long relied on heuristics based on physical insight; however, recent efforts to model conformational ensembles and to make sense of sparse, ambiguous, and noisy data have revealed the value of detailed, quantitative physical models in structure determination. We review these two key challenges, describe different approaches to physical modeling in structure determination, and illustrate several successes and emerging technologies enabled by physical modeling.

**Highlights:**

- Quantitative physical modeling is emerging as a key tool in structure determination
- There are different approaches to incorporate physical modeling into structure determination
- Modeling conformational ensembles and making sense of sparse, noisy, and ambiguous data are two challenges where physical modeling can play a prominent role

## Introduction

Heuristics derived from physical insight have always played an import role in biomolecular structure determination. However, more rigorous quantitative physical models are increasingly used to transform experimental data into structures and ensembles. Physical approaches become more important as the biomolecular system of study becomes more flexible and conformationally heterogeneous (Figure 1), and as experimental data becomes sparse, ambiguous, or noisy (Figure 2). Systems with these characteristics have recently come into focus, due to both the recognition of the importance of conformational heterogeneity and the emerging range of experimental techniques that can provide incomplete information about protein structures [1–5].

**Figure 1:**
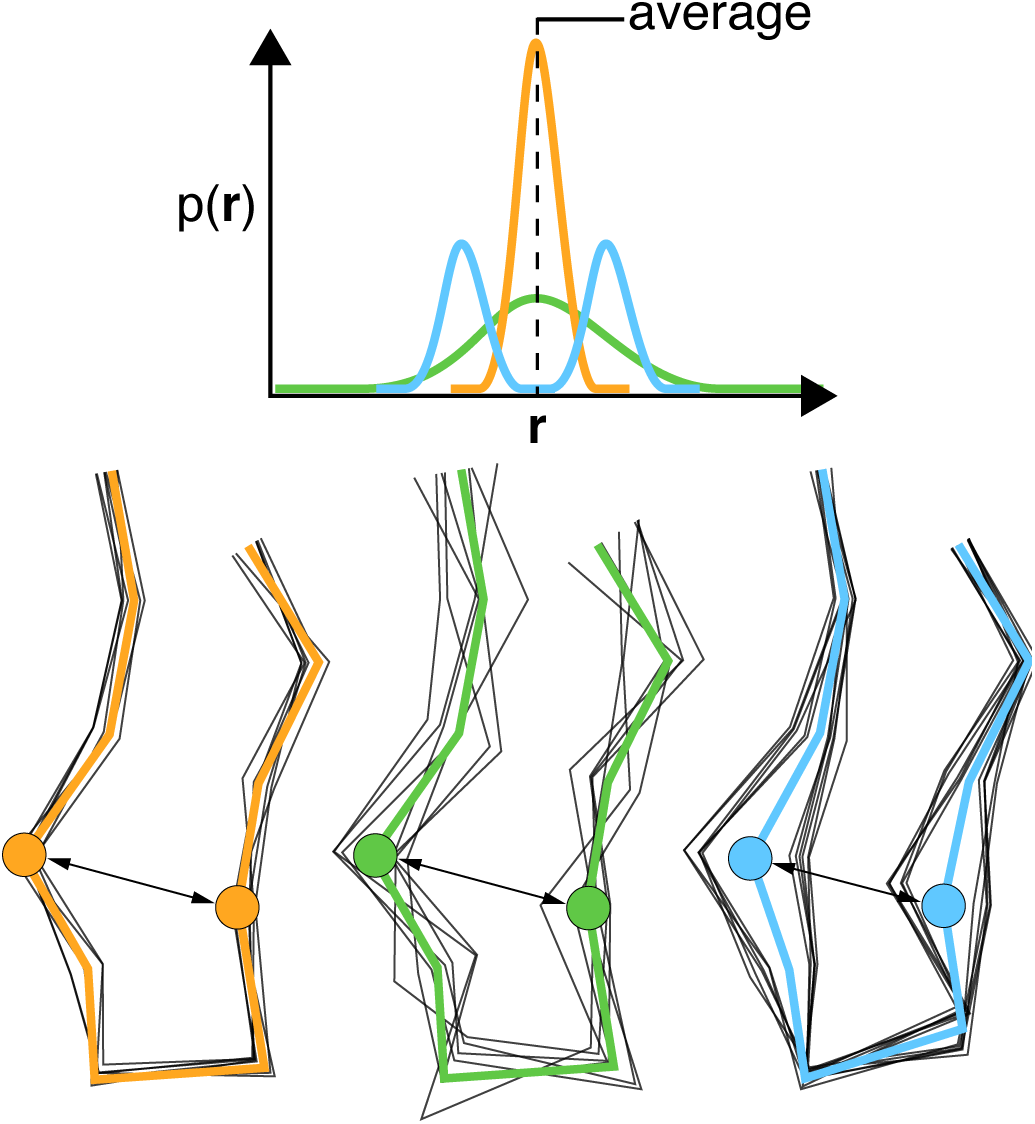
Most experiments measure ensemble averages, which poses a challenge as systems become more flexible, heterogeneous, and dynamic. This figure illustrates a thought experiment, comparing three different ensembles with the same average for some observable, but different conformational ensembles.

**Figure 2:**
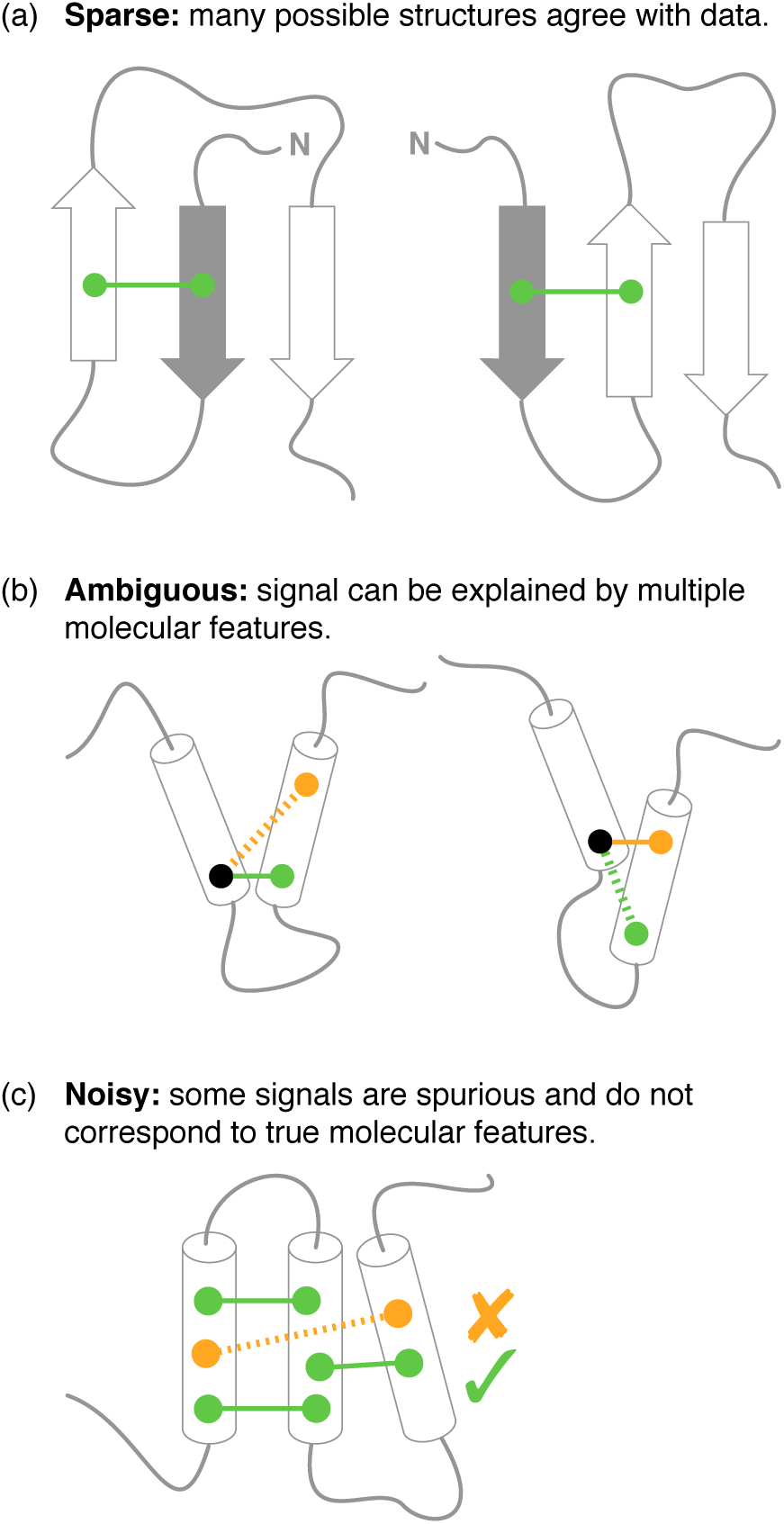
Conceptual illustration of the challenges faced in integrative structural biology and other applications where the data is sparse, ambiguous, and noisy.

Physical modeling has become increasingly powerful over time, driven by improvements in computer power, improved models of energy landscapes [6–8], and improved algorithms for conformational [9–12] and data-driven [13–17] sampling. Combined with advances in experimental methodology, these developments are leading to a new era in structural biology where physical modeling plays a pivotal role [18–20].

In this review, we outline two challenges where physical modeling can make contributions to structure determination, overview some recent successes, and provide a perspective on emerging areas where physical modeling will be important.

### There are several emerging challenges in structural biology

#### Challenge 1: Modeling conformational ensembles

When we refer to “the structure” of a biomolecular system, we are actually referring to some continuous cloud of structures in the neighborhood of a representative structure. While historically the single structure viewpoint has dominated in structural biology, there is increasing recognition of the importance of heterogeneity and dynamics.

Most measurements in structural biology are ensemble averages, where the observed signal comes from the average across many molecules. The challenge of interpreting such averaged data increases as the conformational ensemble becomes more heterogeneous. A simple thought experiment illustrates the central concept (Figure 1), where three systems have the same average for some observable, but different conformational distributions. One system (orange) is tightly clustered, where the average conformation provides an excellent representation of the ensemble. Another system (green) has a broad distribution, where the average conformation is only somewhat representative. The final system (blue) has a multimodal distribution, where the average conformation is improbable and not representative of the underlying ensemble at all. As the experimental average is the same in each case, modeling is critical to making correct inferences about the ensemble.

#### Challenge 2: Making sense of sparse, ambiguous, and noisy data

An increasing variety of experimental methods can provide partial information about the structure of a system [1–5]. While these experiments provide only an incomplete picture, their appeal is that they are often applicable to a wide range of systems, including those where traditional approaches have proven intractable.

Figure 2 shows several common pathologies. First, the data may be sparse, often only providing information about a few degrees of freedom. Second, the data may be ambiguous, where there are multiple molecular features that could explain a particular signal, e.g. an NMR experiment might tell us that two protons are close together, but not specifically which ones. Finally, experimental data is almost always corrupted by noise, which must be interpreted as such to avoid over-fitting. Noise comes in many forms, ranging from simple additive noise (often modeled by an appropriate distribution, e.g. Gaussian noise) to more challenging cases where experimental artifacts lead to the presence of false-positive and false-negative signals.

### What do we mean by physical modeling?

The term “physical modeling” encompasses many approaches, ranging from physically-motivated heuristics to models rooted in rigorous statistical mechanics. Heuristic approaches are motivated by physical considerations and empirical observations. One example is the use of stereochemical restraints during the refinement of X-ray crystal structures [21] that prevent physically impossible bond lengths and overlap between atoms, even though these unrealistic features might lead to näive improvements in the agreement with experimental data. These heuristics are not a comprehensive physical description of biomolecular structure—clearly, one could not hope to predict the correct fold of a protein using only simple stereochemical restraints.

Conversely, statistical mechanics is a rigorous, comprehensive theory that connects the probability 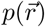 of observing a particular conformation with the potential energy 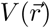 through the Boltzmann distribution:

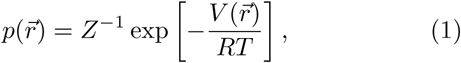

where *R* is the gas constant, *T* is the absolute temperature, and *Z* is a normalization constant called the partition function. Typically, the potential energy is modeled using an empirical approximation called a *force field* [6, 7]. Samples from 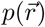 are generated using molecular dynamics or Monte Carlo simulations, often augmented by various enhanced sampling algorithms [10, 12, 13, 22].

Rosetta is another example of physical modeling [8]. Although the underlying philosophy and parameterization of Rosetta differ substantially from those of statistical mechanical models, the underlying goal is essentially the same—to reproduce the conformational landscape of a biomolecular system of interest.

### There are different approaches to incorporating physical models into structure determination

Constructing structural models of a biomolecular system from one or more experimental datasets can be cast as an inference problem that can be solved through a variety of approaches (reviewed in [19, 20]). However, the distinctions between these approaches can often be subtle. In this section, we outline several characteristics that distinguish different approaches.

#### Characteristic 1: What is the nature of the likelihood function?

The likelihood, *ℒ*(*θ |D*) ∼ ℙ(*D |θ*), is central to many methods, where *D* is the observed data and *θ* is a set of parameters specifying the structural ensemble, e.g. atomic coordinates. The likelihood is typically built by combining a *forward model*, which calculates the experimental observable from a given structure or ensemble, with a *noise model*, which calculates the probability of a given deviation from the measured value [23, 24]. Success depends critically on the quality of the forward model, as any inaccuracies lead directly to errors in the final ensemble.

One must distinguish between likelihoods that consider single structures from those that consider ensembles. We refer to these as *single-structure* and *ensemble* likelihoods; they are also referred to as likelihood *functions* and *functionals*, respectively. Use of a single-structure likelihood will make all members of the ensemble match experiment, which may be acceptable for systems with only limited heterogeneity. But ensemble likelihoods, which ensure that model averages match observations, are essential for more heterogeneous systems.

The relationship between structure and observable may be non-linear. For example, NOE and FRET measurements are proportional to ⟨*r*^−6^⟩, leading to averages dominated by (potentially) rare short-distance conformers. Similarly, observation of chemical cross-linking indicates occasional proximity of two residues, but as the cross-links are irreversible once made, one cannot relate the average degree of cross-linking to the average distance between the residues. Furthermore, rather than only providing averages, some experiments, like EPR or single-molecule FRET, can provide *distributions*, which are potentially far richer. In all cases, the likelihood function must correctly capture these relationships in order to avoid biasing the calculated ensemble.

#### Characteristic 2: What statistical formalism is used?

Maximum likelihood (ML) and Bayesian statistical approaches are two common ways to solve the structural inference problem. Maximum likelihood methods aim to find the single best set of parameters, 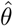, that maximize the likelihood function. Näive ML methods determine the parameters based entirely on the data, making these methods sensitive to noise and notoriously prone to over-fitting. To mitigate this, it is common to use penalized ML methods, where the aim is to minimize:

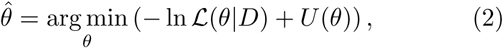

where *U* (*θ*) is a penalty function that may include simple restraints on stereochemistry, more detailed energetic models given by force fields, or *ad hoc* penalty terms motivated by physical considerations, e.g. the use of restraints on crystallographic B-factors [25]. Such penalty terms are a form of regularization and ensure individual conformations are physically reasonable, preventing over-fitting.

The Bayesian approach offers a different perspective [23, 24], where one seeks to find the joint distribution of parameters given the data. Bayes theorem is a simple and elegant statement,

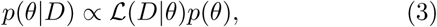

that combines prior understanding with new information in a statistically consistent way. The quantity of interest is the posterior distribution, *p*(*θ |D*), which is obtained by combining the likelihood with the prior, *p*(*θ*). The prior, often modeled as the Boltzmann distribution, plays a similar role to the penalty terms in ML methods, in that it encodes our knowledge about what structures are *a priori* probable.

A major differences between the two approaches is that ML methods provide a *point estimate* of the best parameters, whereas Bayesian methods produce a *joint distribution* of parameters—or, more commonly, a set of samples from the joint distribution generated by molecular dynamics or Monte Carlo sampling. Bayesian methods naturally provide an estimate of uncertainty that also captures the coupling between different parameters. Bayesian maximum *a posteriori* (MAP) estimators seek to find the mode of the posterior distribution, i.e. the single most likely set of parameters, blurring the lines between ML and Bayesian approaches.

Bayesian approaches readily allow for the inclusion of “nuisance parameters", which are unobserved, but nevertheless influence the results of inference. Examples include the true values of experimental observables (opposed to noisy observations) or the exact values of empirical constants used in forward functions. The Bayesian approach [23, 24] is to treat these as nuisance parameters, with appropriate priors, that are inferred jointly with the rest of the model. The resulting distribution gives the likely values of the nuisance parameters and their influence on the generated ensemble.

Many approaches used in structural biology do not neatly fall into either the ML or Bayesian frameworks. These are often more *ad hoc* combinations of physically-motivated scoring functions and sampling strategies that do not produce a well-defined ensemble. For example, traditional NMR refinement generates collections of low-energy structures, but these do not correspond to welldefined statistical or thermodynamic ensembles. Although these methods have less rigorous statistical underpinning, they are very common in structural biology and obviously quite useful.

#### Characteristic 3: What principle is used to regularize ensembles?

The previous section outlined how penalty terms or priors can be used to regularize *individual structures*. However, without additional regularization, *ensemble models* become prone to over-fitting due to poor data-to-parameter ratios. For example, it is uncommon to see multi-copy refinement of X-ray crystal structures [26]. Phillips and co-workers undertook a systematic study of 50 experimental structures, and found that adding up to, on average, *∼* 10 copies yielded improved models [27], with over-fitting occurring beyond that. *Ensemble regularization* methods can help avoid such over-fitting.

One approach is to use the *principle of maximum parsimony*, which seeks to find a minimum representative set of conformations that adequately explain the experimental data. These methods are typically based on re-weighting or selection. A pool of conformations is generated from a prior distribution using Monte Carlo or molecular dynamics sampling. Weights are then assigned to each conformation to bring calculated averages into agreement with experimental observations. A variety of approaches are possible [28–35] with several ways to choose the number of representative conformations, including user specification [29–31], clustering [32, 33], and penalty terms or priors that favor sparse models where most weights are zero [35].

Another approach to ensemble regularization is based on the *principle of maximum entropy* (MaxEnt), which posits that the distribution that best reflects our current state of knowledge is one that agrees with experimental observations, while simultaneously maximizing entropy [36, 37]. The Boltzmann distribution (Eq. 1) is the MaxEnt distribution over conformations, subject to a constraint relating the average energy and temperature [37]. When experimental observations are included, one arrives at [38]

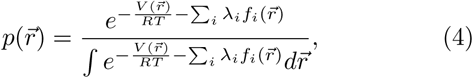

where 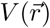 is the potential energy, and {*λ*_*i*_} are Lagrange multipliers that must be determined in order satisfy agreement between model averages and observations:

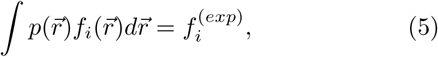

where 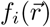 is the function computing the *i*th observable and 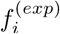 is the corresponding experimental observation. Equivalently, one can minimize the Kullback-Leibler divergence between the inferred ensemble and the Boltzmann distribution, while satisfying the constraints of Eq. 5.

MaxEnt methods fit the Lagrange multipliers, rather than fitting the conformations directly. Because there is one Lagrange multiplier for each observation, the data to parameter ratio remains constant as the number of conformations increases, which allows for large ensembles without over-fitting. The magnitude of the Lagrange multipliers provides insight into how strongly the prior was perturbed to match each experimental observation.

A variety of methods exist for MaxEnt ensemble determination, as recently reviewed in [19, 20]. Approaches include re-weighting [39, 40], iterative schemes [38, 41–43], time-dependent potentials [44, 45], and restrained ensemble or replica-averaged schemes [40, 46–52]. Pitera and Chodera [38] identified a link between Eq. 4 and restrained ensemble schemes, which has been further clarified [40, 52, 53].

Although potentially less prone to over-fitting, care is still required when using maximum parsimony and MaxEnt methods. These approaches are sensitive to the quality of the prior distribution and can be expected to perform poorly when the prior distribution differs substantially from the true distribution.

### The term “ensemble” is highly overloaded in structural biology

In statistical mechanics, the term “ensemble” has a specific technical meaning: the probability distribution over all possible conformations of the system under specified conditions. Unfortunately, in structural biology it has become common to refer to almost any collection of conformations as an ensemble, which can be confusing.

There are several reasons that an ensemble may be heterogeneous. First, the ensemble may be a *thermodynamic ensemble*, where the heterogeneity is intended to reflect the true physical heterogeneity of the system at equilibrium. Second, the ensemble may be an *uncertainty ensemble*, where the heterogeneity reflects that there may be many possible structures compatible with sparse, ambiguous, or noisy data. Third, one can also distinguish between uncertainty ensembles composed of samples from well-defined statistical distributions, e.g. from Bayesian approaches, and those that are simply collections of structures generated by some procedure. Caution is required when interpreting uncertainty ensembles, as the heterogeneity present is not necessarily predictive of the true heterogeneity.

Whenever one interprets an “ensemble” there are several key characteristics that must be considered. Does the ensemble come from a maximum likelihood or Bayesian approach, or from some other method? Is a single-structure or ensemble likelihood used? Do the structures sampled come from a well defined distribution? How are errors modeled? What priors or penalty functions are used?

### Physical modeling offers solutions to challenges in structural biology

Hummer and co-workers introduced a Bayesian ensemble refinement method BioEN, a combination of replica ensemble refinement and the Ensemble Refinement of SAXS (EROS) method, combining the principles of both restraining and reweighing [40].

Ensemble heterogeneity explains much of the difficulty in characterizing intrinsically disordered proteins (IDPs) experimentally, as they are ensembles of inter-converting conformations [54, 55]. The Bayesian weighting method is an approach for characterizing an ensemble of IDPs where the weights are defined using a Bayesian estimate from calculated chemical shift data [33]. This method has been successful in determining the relative fractions of mutated structures in an ensemble for aggregative proteins [56].

Of all biomolecules, RNA has been perhaps the most challenging to simulate with molecular dynamics, as current force fields are not accurate enough to reproduce experiments [45]. Cesari and co-workers used a MaxEnt approach to bias RNA simulations to match J-coupling experiments [45] and demonstrate that this can be used to develop a self-consistent, transferable force field correction.

High ambiguity driven biomolecular docking (HAD-DOCK), is a data-driven docking approach, that can take highly ambiguous data from different sources and convert them into distance restraints to guide docking processes [57, 58]. HADDOCK has been used to study protein complex interfaces using cryo-EM data [59] and protein ligand complexes using sparse intermolecular NOEs [60].

The Integrative Modeling Platform (IMP), is a flexible software suite aimed at integrative structural biology, which facilitates development of integrative applications, models and methods, and allows incorporation of data from diverse sources [15]. Large protein complex structures have been modeled with IMP using *in vivo* FRET data through a Bayesian approach [61], and using a combination of cross-linking data with biochemical and EM localization data [62].

Rosetta is an extensive software suite aimed at protein structure prediction and molecular design. There are several applications of Rosetta with sparse experimental data, where Monte Carlo-based fragment assembly is guided towards native structures by data [63]. Backbone chemical shifts and distance restraints have been used to guide structure determination [64]. Also, paramagnetic relaxation enhancement (PRE) [65], pseudo-contact shift (PCS) [66], and residual dipolar coupling (RDC) [67] restraints have been used to similar effect. Recently, the RASREC algorithm was developed, which yields better models with narrower sampling [17, 68] and has been applied to NMR on deuterated samples up to 40 kDa [69, 70].

Metainference, a recent approach based on Bayesian inference, can address statistical and systematic errors in data produced by high-throughput techniques, and can handle experimental data averaged over multiple states [14]. It is suitable for studying structural heterogeneity in complex macromolecular systems. A combination of Metainference and Parallel-bias Metadynamics (PBMetaD), an accelerated sampling technique, provides an efficient way of simultaneously treating error and sampling configuration space in all-atom simulations [9]. Coupling Metainference and Metadynamics has been particularly successful in characterizing structural ensembles of disordered peptides [71, 72].

Modeling Employing Limited Data (MELD) is a Bayesian approach that combines statistical mechanics, detailed allatom physical models [7], and enhanced sampling to infer protein structures from sparse, ambiguous, and noisy data [13]. MELD was specifically designed to be robust in the presence of false-positive signals, and has been applied to EPR, NMR, and evolutionary data [13], *de novo* prediction of protein structures based on simple heuristics [73, 74], and mutagenesis guided peptide-protein docking [75, 76].

### Physical modeling is enabling emerging techniques in structural biology

Advances in physical modeling will be key to enabling technologies for new approaches to structure determination. Below we outline just a few—of many—emerging techniques where the ability to model ensembles and to successfully treat sparse, ambiguous, and noisy data will be critical.

Chemical cross-linking detected by mass spectrometry is emerging as a potentially powerful tool in structure determination. Developments have focused on improvements in instrumentation [4, 77], cross-linking chemistries [78–80], and data analysis [78, 79, 81, 82]. These techniques are extremely sensitive, but the data can be highly ambiguous, both false-positive and false-negative signals are common, and one cannot relate the degree of cross-linking to the average distance. Such data has recently been used as restraints to guide Monte Carlo [83], molecular dynamics [84, 85], and integrative modeling [81, 82] approaches. The use of cross-linking restraints for structure prediction was recently assessed during the 11th round of Critical Assessment of Structure Prediction [86, 87] and various shortcomings—both in experiment and modeling— were identified.

X-ray diffuse scattering experiments can produce information about correlated motions in proteins that is complementary to the information obtained from the more typically analyzed Bragg scattering [88, 89]. Wall and co-workers found good agreement between long molecular dynamics simulations and measured diffuse scattering [89], even in the absence of any fitting. The development of suitable ensemble refinement schemes would bring the models into even better agreement with experiment and would provide a powerful new tool for studying correlated motions of proteins.

Recent work has demonstrated the utility of paramagnetic relaxation enhancement measurements in solid-state NMR [90, 91]. These experiments provide less structural information than traditional protein NMR experiments, but, combined with suitable computational modeling, represent an increasingly viable avenue for structure determination [65, 91].

Finally, recent work has demonstrated the possibility of inferring residue–residue contacts from coevolution analysis of homologous sequences [92–94], commonly referred to as evolutionary couplings. Baker and co-workers were recently able to create models for 614 protein families with unknown structures [95], several of which had folds that are not in the Protein Data Bank. Montelione and co-workers combined evolutionary couplings with sparse NMR data, which provide complementary restraints for modeling, to correctly determine structures for proteins up to 41 kDa [3].

### Conclusion and future perspectives

Physical insight has always been integral to structural biology, but the dual challenges of modeling ensembles and making sense of sparse, ambiguous, and noisy data mean that quantitative physical models will become an increasingly important part of modern structural biology. How can we avoid misinterpreting measurement noise as structural heterogeneity? How can we recognize rare, but important, conformations buried within noisy data? Making progress requires: (1) better experiments, including those that are sensitive for rare conformations [96, 97] or provide distributions [97–99]; (2) improved models of the noise and error inherent in the experimental data; (3) accurate methods to back calculate observables from structural ensembles; (4) accurate physical models that can correctly reproduce the conformational landscapes of biomolecules; and (5) suitable statistical frameworks to interpret all of this information in a coherent fashion. In the future, we anticipate that approaches combining statistical inference, physical modeling, and experiments will allow us to better understand the dynamic and heterogeneous nature of conformational ensembles, even in the presence of noisy, sparse, or ambiguous data, which will be key to addressing important biological questions.

### Important References

Papers of interest have been highlighted as:

* of special interest

** of outstanding interest

** [20] A review on approaches that combine experimental and computational methods to determine structural ensembles of dynamic proteins.

**[19] A concise review on maximum entropy approaches. The authors highlighted three papers which explored an important link between replica-averaged ensemble refinement principle and maximum entropy method.

**[18] An important perspective on the relationship between experimental data and computational techniques, and the role of integrative structural biology.

**[23] A key paper on Bayesian inference, defining the commonly applied inferential structural determination methodology and indicating the importance of developing probabilistic methods for structure determination.

**[38] This paper makes use of maximum entropy methods to develop ensemble-averaged restraints for biasing molecular simulations, noting the success of a physics-based approach compared to other refinement schemes.

*[52] This paper demonstrates the statistical equivalence of principle of restrained-ensemble simulations and the the maximum entropy approach.

*[53] This paper justifies the use of the maximum entropy approach to define experimental data-driven restraints for simulations and demonstrates its equivalence to replicaensemble simulations.

*[14] This paper introduces a Bayesian inference method to account for different sources of error in experimental data in modeling structural ensembles of complex macromolecular systems.

*[15] This paper introduces the new and developing Integrative Modeling Platform (IMP) software package. The authors highlight its flexible capability to incorporate a variety of experimental data, and to generate and develop new models and representations.

*[13] This paper describes Modeling Employing Limited Data (MELD), highlighting its unique Bayesian methodology for determining protein structure, and demonstrating its ability to incorporate a variety of experimental data.

*[70] This paper introduces the RASREC Rosetta approach, describing its improvements over regular CS-Rosetta in detail, and exhibiting its capability to develop models closer to the native structure.

## Acknowledgements

This work was supported by funding from the Natural Sciences and Engineering Research Council of Canada. JLM is a Tier 2 Canada Research Chair.

